# The mitochondrial genome of *Oryza sativa* L. cv. Taichung 65 (Poaceae)

**DOI:** 10.1101/2022.03.02.482735

**Authors:** Hiroyuki Ichida, Tomohiko Kazama, Shin-ichi Arimura, Kinya Toriyama

## Abstract

A complete mitochondrial genome sequence was determined by a hybrid approach with long- and short-read sequences in a *japonica* rice cultivar Taichung 65. The assembled mitochondrial genome was 401,060 bases in length and an overall GC content of 43.88% and predicted to harbor 57 protein-encoding, 18 transfer RNA, and 3 ribosomal RNA genes. Phylogenetic and sequence polymorphism analysis indicated that Taichung 65 is classified as typical *japonica*. The mitochondrial genome resource would serve as a fundamental tool to investigate cytoplasmic male sterility in rice.

## Main text

Taichung 65 is a *japonica* rice cultivar *(Oryza sativa* L.) reportedly bred by crossing two old Japanese landraces, Kameji and Shinriki. The cultivar has been predominantly used as a nuclear donor plant for the production of alloplasmic cytoplasmic male sterile (CMS) lines in Japan (Toriyama, 2021). For studies of such alloplasmic CMS lines, the Taichung 65 mitochondrial genome nucleotide sequence is necessary as a positive control. In this study we determined the whole genome sequence of the Taichung 65 mitochondria.

Seeds of Taichung 65 were obtained from Ryukyu University as a maintainer line for Chinsurah Boro II-Taichung 65 type CMS (Shinjyo, 1969). The cultivar was maintained by self-pollination in Tohoku University. Genomic DNA was extracted from fully expanded leaf blades using the Genomic-tip 100 (Qiagen, Hilden, Germany) in accordance with the manufacturer’s protocol. Rice mitochondria were purified by density centrifugation, following methods described elsewhere (Tanaka *et al.,* 2004, Kazama *et al.,* 2008). Our detailed protocol for mitochondrial isolation is presented in supplemental data. Mitochondrial DNA was extracted using a DNeasy Plant Mini Kit (Qiagen, Hilden, Germany). The purified mitochondrial DNA was sequenced with a SMRT cell on a PacBio RS II sequencer at Macrogen (Seoul, Korea). A total of 680,369,198 bases spanning 79,106 filtered subreads was obtained. The mean subread length and N50 of the subreads were 8,600 and 11,976 bases, respectively. Bioinformatics analysis was performed on the Hokusai massively parallel computing system, operated by Information Systems Division, RIKEN, under project number Q21208. *De novo* assembly was performed using Canu (version 2.1.1; (Koren *et al.,* 2017). To optimize assembly conditions, we scanned minimal subread lengths between 5,000 and 20,000 bases, in increments of 2,500 bases, as well as minimum overlap lengths between 500 and 5,000 bases, in increments of 500 bases. Based on total contig counts and N50 values, we ascertained that a minimal subread length of 7,500 bases with an overlap length of 1,000 bases gives a good balance between contiguity (total number of contigs) and coverage (total length). The contig sequences were polished for better sequence accuracy with Pilon (version 1.24; (Walker *et al.,* 2014) and the UnifiedGenotyper and

FastaAlternateReferenceMaker tools in GATK (version 3.8.1.0; (McKenna *et al.,* 2010) using short-read sequences from Taichung 65 available from the Sequence Read Archive (SRA; BioProject PRJNA280553). The final polished contigs were assembled by manually inspecting similarity with the published mitochondrial genome sequence from rice cultivar Nipponbare (GenBank accession code DQ167400). This resulted in a single mitochondrial genome sequence. The annotated mitochondrial genome sequence is available under the accession code LC697740 in DDBJ/GenBank. The high-throughput sequencing results were deposited in the SRA under BioProject PRJNA808844.

The complete mitochondrial genome consists of a single, non-circular sequence with a total length of 401,060 bases and an overall GC content of 43.88%. The total length of the Taichung 65 mitochondrion is approximately 20% shorter than that of Nipponbare (490,669 bases in length) but has a similar GC content (43.85%). Two major structural differences between the two mitochondria genome sequences are apparent: an inversion between positions 120,337 and 182,819 (in Taichung 65; 145,000 and 209,999 in Nipponbare) and a missing duplication between positions 254,955 and 302,871 (in Taichung 65; 420,000 to 469,999 in Nipponbare). The existence of these structural variations was verified by manually inspecting the aligned reads from long- and short-read sequencing. Some mapped reads support the inversion and missing duplication; however, reads supporting the Nipponbare-type structure were also observed in both cases. This indicates that both types of structures exist in the Taichung 65 mitochondria, presumably due to spontaneous flipping by endogenous recombination.

Gene prediction from the assembled Taichung 65 mitochondrial genome sequence indicates that all tRNA genes and commonly found mitochondrial genes are conserved in the assembled sequence. The examination of polymorphisms of segmental sequence variations at 11 loci have been reported to discriminate between typical *japonica* and *indica* rice lines (Luan *et al.,* 2013). In our case, all of the corresponding Taichung 65 polymorphisms coincide with those of Nipponbare, confirming that the Taichung 65 mitochondrial genome should be classified as typical *japonica*. A phylogenetic tree was constructed based on the 38 conserved protein-coding genes (Figure 1). The result confirmed that Taichung 65 is most closely related to *O. sativa* ssp. *japonica* cultivars, although both subspecies in *O. sativa,* as well as its wild-relative *O. rufipogon,* are close together in mitochondrial protein level. The Taichung 65 mitochondrial genome sequence will now be available as a fundamental resource for the investigation of various molecular mechanisms defined by mitochondria. Further analysis of additional rice cultivars and wild relatives should elucidate the origin and evolution of such mechanisms.

**Figure 1.**
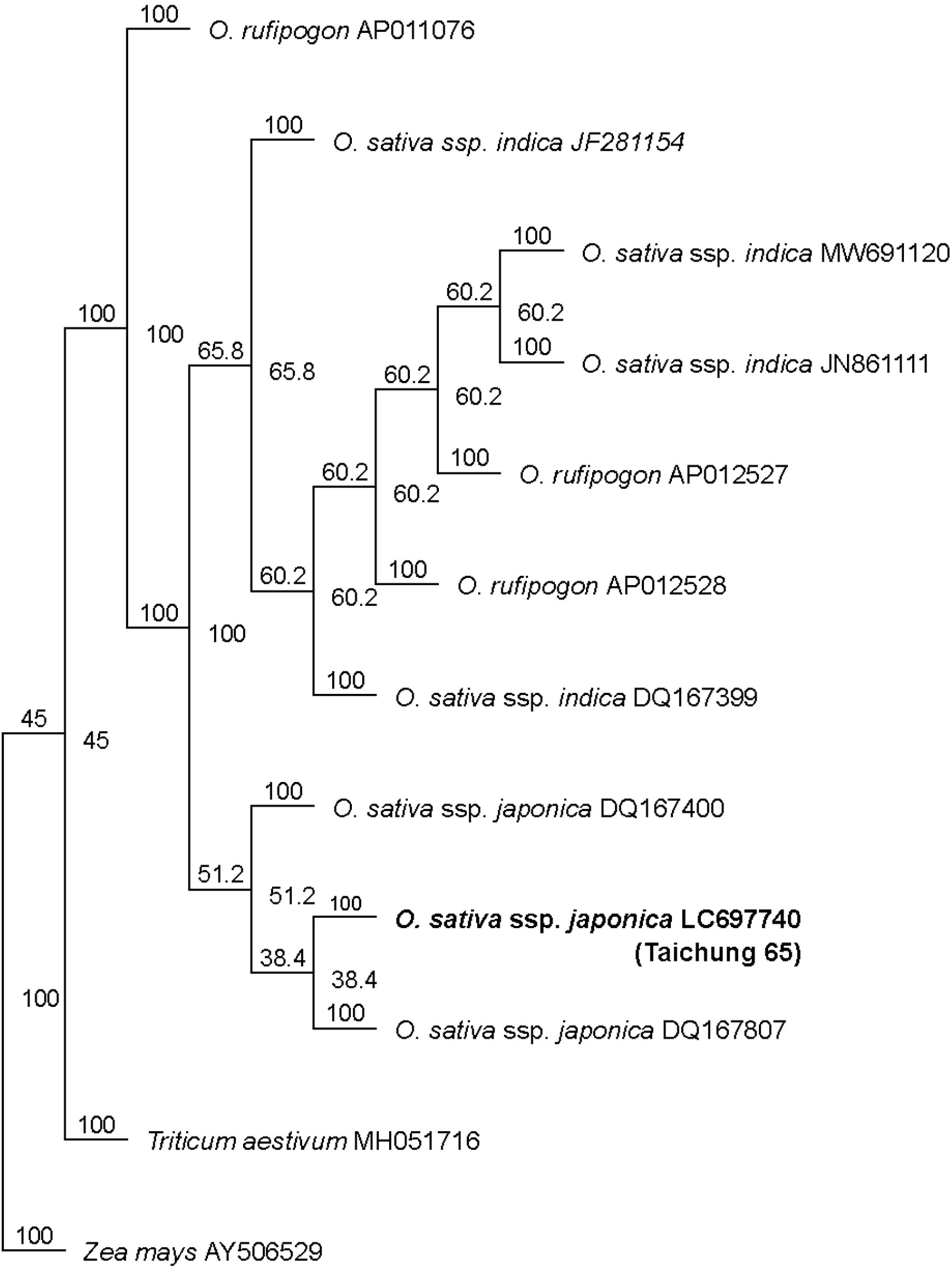
A UPGMA tree for the Taichung 65 (LC697740) and other representative GenBank records in *Oryza* spp. The tree was constructed using 38 conserved protein-coding genes. The tree was constructed with the Geneious Tree Builder tool with the Jukes-Cantor distance model. The numbers at the nodes are bootstrap percent probability based on 1000 replicates.

## Data availability

The data that support the reported study is available in DDBJ/GenBank with the accession code LC697740. The high-throughput sequencing results were deposited in Sequence Read Archive under BioProject PRJNA808844.

